# Unified single-cell analysis of testis gene regulation and pathology in 5 mouse strains

**DOI:** 10.1101/393769

**Authors:** Min Jung, Daniel Wells, Jannette Rusch, Suhaira Ahmed, Jonathan Marchini, Simon Myers, Donald F. Conrad

## Abstract

By removing the confounding factor of cellular heterogeneity, single cell genomics can revolutionize the study of development and disease, but methods are needed to simplify comparison among individuals. To develop such a framework, we assayed the transcriptome in 62,600 single cells from the testes of wildtype mice, and mice with gonadal defects due to disruption of the genes *Mlh3*, *Hormad1*, *Cul4a* or *Cnp*. The resulting expression atlas of distinct cell clusters revealed novel markers and new insights into testis gene regulation. By jointly analysing mutant and wildtype cells using a model-based factor analysis method, SDA, we decomposed our data into 46 components that identify novel meiotic gene regulatory programmes, mutant-specific pathological processes, and technical effects. Moreover, we identify, *de novo*, DNA sequence motifs associated with each component, and show that SDA can be used to impute expression values from single cell data. Analysis of SDA components also led us to identify a rare population of macrophages within the seminiferous tubules of *Mlh3-/-* and *Hormad1-/-* testes, an area typically associated with immune privilege. We provide a web application to enable interactive exploration of testis gene expression and components at http://www.stats.ox.ac.uk/~wells/testisAtlas.html

## Introduction

The testis is an amalgamation of somatic cells and germ cells that coordinate a complex set of cellular interactions within the gonad, and between the gonad and the rest of the organism (**Figure 1A**). The key function of the testis is to execute spermatogenesis, a developmental process that operates continually in all adult mammals. The mechanisms of this process are important for the evolution, fertility and speciation of all sexually reproducing organisms.

**Figure 1.**
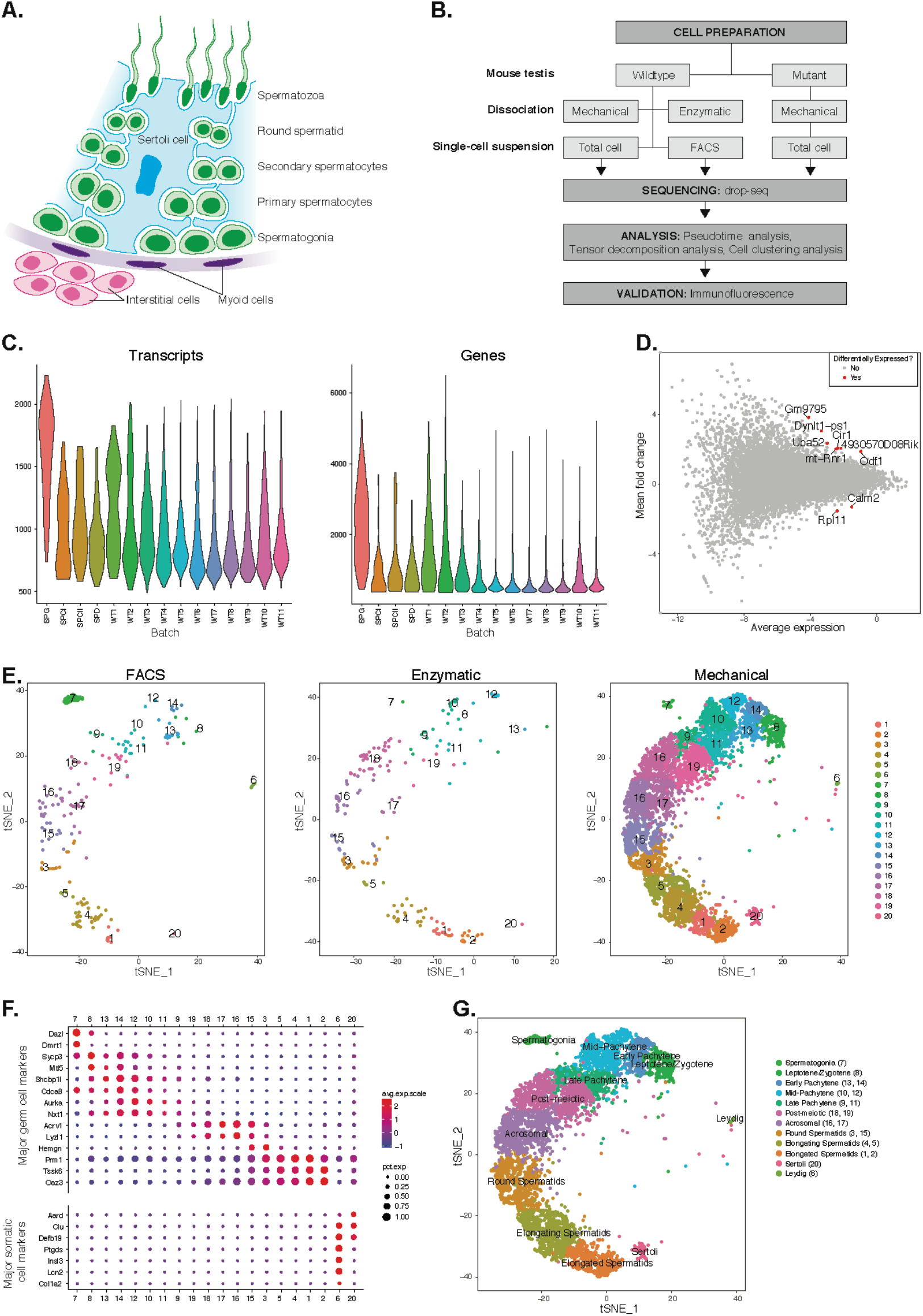
Mapping cellular diversity in the adult testis using single-cell expression profiling. (A) Anatomy of the testis. Adult testis are comprised of germ cells (spermatogonia, primary spermatocytes, secondary spermatocytes, spermatids and spermatozoa) and somatic cells. Within the seminiferous tubules, there is a single population of somatic cells (Sertoli). The tubules are wrapped by muscle-like “myoid” cells. Outside the tubules are a heterogeneous, poorly defined population of “interstitial” somatic cells. (B) Overview of the experiments. To establish the utility of single-cell profiling for testis phenotyping, we performed a series of experiments (i) comparing the quality of traditional enzymatic dissociation and more rapid mechanical dissociation, (ii) comparing the expression profiles of cells from total testis dissociation to testicular cells of known identity purified by FACS, (iii) comparing expression profiles of wildtype animals to cells isolated from 4 mutant strains with testis phenotypes. (C) We used Drop-seq to profile 26,615 cells from wildtype animals, with an average of 1,032 transcripts/cell and 929 genes/cell. We compared chemical dissociation (SPG, WT1, WT2) and mechanical dissociation protocols (all others). (D) Expression profiles from enzymatic and mechanical dissociation showed excellent concordance (R=0.95). (E) We used t-SNE to visualize k-means clustering of 19,153 cells in 12 clusters. No obvious batch effects were detected when comparing the t-SNE clustering location of cells isolated by FACS, or either of the two total testis dissociation protocols. (F) Expression levels of known germ cell and somatic cell markers were used to assign labels to clusters. (G) Label assignment clearly indicates a spatial organization of testis cells in t-SNE space, with one dimension (y-axis) clearly reflecting the continuous ordering of germ cells along the developmental trajectory of spermatogenesis, with somatic cell populations flanking the germ cells in small pockets.

A deeper understanding of the transcriptional programme of spermatogenesis has potential applications in contraception ^1^, *in vitro* sperm production for research and the treatment of infertility ^2^, and the diagnosis of infertility, among others. Yet, the study of the highly dynamic transcriptional programmes underlying sperm production has previously been limited by the cellular complexity of the testis. It is comprised of at least 7 somatic cells types, and at least 26 distinct germ cell classes that identifiable by morphology ^3^.

The testis has a number of unique genomic features: its transcriptome has by far the largest number of tissue specific gene (over twice as many as the 2^nd^ ranked tissue the cerebral cortex – with which the testis shares an unusual similarity) ^4,56^; it contains the only cells in the male body with sex chromosome inactivation; and it features dramatic chromatin remodelling, when the majority of histones are stripped away during spermiogenesis and replaced with small, highly basic proteins known as protamines.

Use of genetic tools has also enabled the dissection of the homeostatic mechanisms that regulate spermatogenesis, revealing both cell autonomous and non-autonomous mechanisms. However, most perturbations that disrupt spermatogenesis also change the cellular composition of the testis, frustrating the use of high throughput genomic technologies in the study of gonadal defects. By removing heterogeneity as a confounding factor, single cell RNA sequencing (scRNA-seq) promises to revolutionize the study of testis biology. Likewise, it will completely change the way that human testis defects are diagnosed clinically, where testis biopsy is the standard of care for severe cases.

Here, we performed scRNA-seq on 62,600 testicular cells from the mouse testis, using wild-type animals and 4 mutant lines with defects in sperm production (**Figure 1B**). On the basis of these data, we set out to develop an analysis approach that would allow us to rapidly extract mechanistic insights from joint interrogation of multiple mouse strains; to revisit fundamental questions in spermatogenesis using the new resolution of single-cell analysis; and to establish the utility of scRNA-sequencing for dissecting testis gene regulation in both normal and pathological states.

## Mapping the cellular diversity of the testis

We first tested two methods for testis dissociation: enzymatic dissociation, a slow 2-hour protocol, vs. a rapid 30-minute protocol based on mechanical disruption.^7^ Single cell expression profiles from the two methods showed excellent agreement (r=0.95), and no important differences in cell quality or ascertainment (**Figures 1D-E**); thus we applied the mechanical dissociation approach for the vast majority of the experiments (**Table S1**). We performed scRNA-seq to generate 25,423 cell profiles isolated from total testis dissociations of 11 wild-type animals (WT1-WT11). We compared these to reference data for 296 spermatogonia, 199 primary spermatocytes, 398 secondary spermatocytes, and 299 spermatids, all purified by FACS (**Methods**).

The yield of transcripts per cell was consistent with previous studies on different cell types (**Table S2**) and after QC, we retained 19,153 cells for clustering. Initially, we performed cell clustering. This identified 12 distinct populations, which we visualized using t-distributed Stochastic Neighborhood Embedding (t-SNE) (**Methods**, **Figure 1E**). By inspecting the expression levels of known cell type markers and comparing to FACS-sorted cells, we could unambiguously resolve these populations into 9 subtypes of germ cells and 2 somatic cell populations – Leydig cells and Sertoli cells (**Figures 1F-G**). Along with the expected patterns of expression for known markers, we identified numerous novel markers for all populations, which validated well with immunohistochemistry (**Figure 2A**, **Figure S1**). Noteworthy are the identification of *Kif5b* as an excellent marker of Sertoli cells that provides more extensive coverage of the cell body than the conventional markers *TUBB* and *Vim*, and the identification of *Abhd5* as the first marker specific to the subcellular structure of developing germ cells known as the residual body.

**Figure 2.**
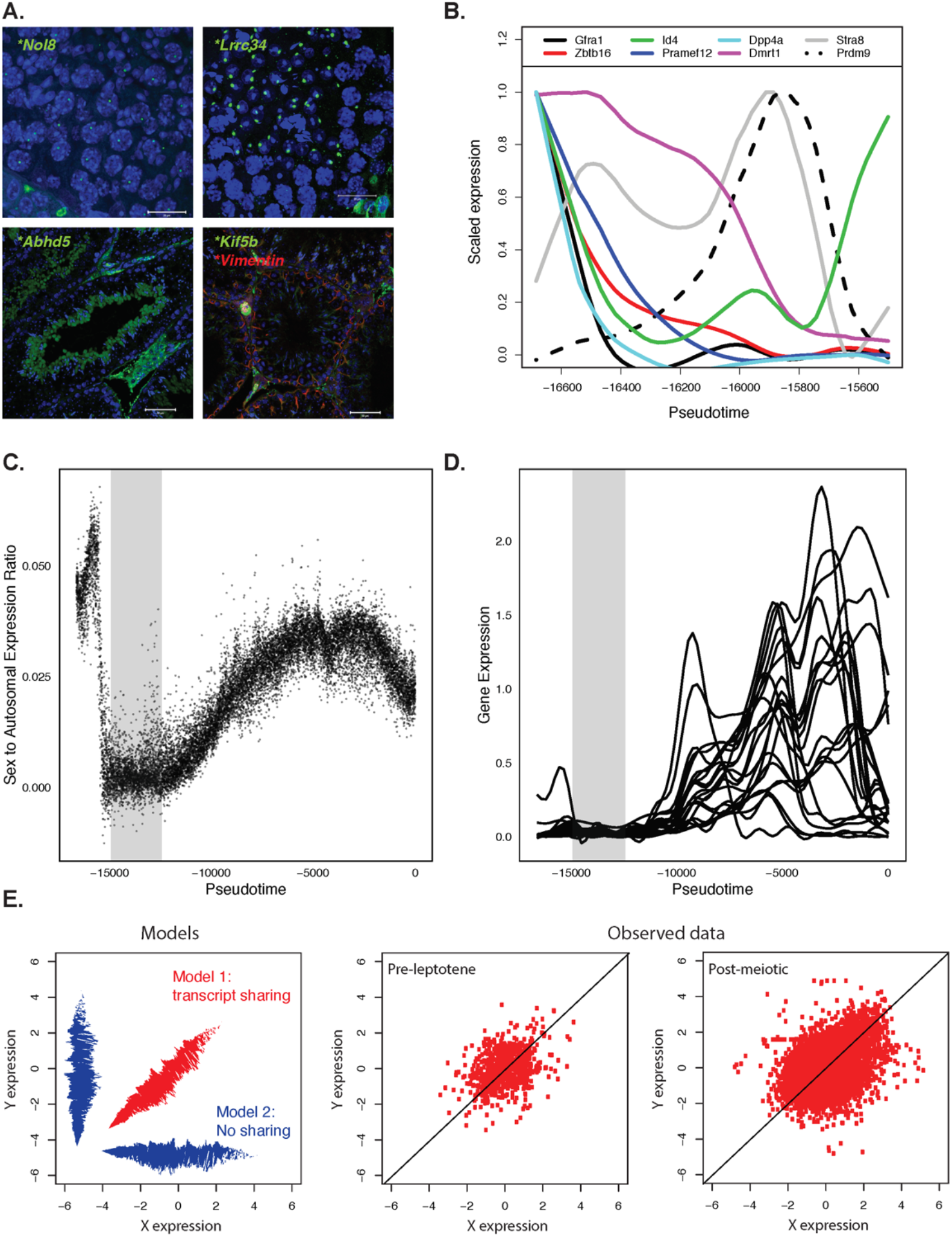
Novel insights from single cell sequencing of wildtype cells. (A) Across all major cell clusters, we identified 28 highly specific gene expression markers that were not previously reported in the literature (**Figure S1**). We attempted to validate 10 of these at the protein level. These included *Nol8* (primary spermatocytes); *Lrrc34* (spermatids); *Abhd5* (residual body); *Kif5b* (Sertoli). (B) Pseudotime analysis provides insights into the expression properties of spermatogonial stem cells. By assessing the expression dynamics of genes that are proposed to act as spermatogonial “stem cell” (SSC) markers (*Id4*, *Gfra1*, and *Zbtb16*) we found that the population at the earliest stages of germ cell development expresses both *Id4* and *Gfra1*. We identified a number of genes with similar expression dynamics to *Id4* and *Gfra1*, but whose role in germ cell biology is poorly studied, including genes *Dpp4a* and *Pramef12*, (main text, **Table 1, Figure S2**). (C) Pseudotime analysis also allow provides quantitative, high-resolution insights into meiotic sex chromosome inactivation (MSCI). Chromosome expression relative to autosomal drops to almost 0 showing near-completeMSCI before gradually partially recovering. (D) No evidence supporting prior report of genes escaping MSCI (E) Competing models of transcript sharing in haploid spermatocytes, compared to the observed pattern, which clearly demonstrate extensive sharing of transcripts from the sex chromosomes.

Single cell RNA sequencing provides new opportunities to assess important open questions in the field of spermatogenesis. One long-standing challenge has been to identify molecular markers of spermatogonial stem cells (SSCs), operationally defined as cells which can successfully reconstitute spermatogenesis after transplantation into an empty niche. Various genes have been nominated as expression markers of spermatogonia stem cells, including *Zbtb16* (aka *Plzf*) ^8^, *Id4 ^9^*, *Gfra1 ^10^*, *Nanos3 ^11^*, and *Lin28a ^12^*. We performed pseudotime modeling (Methods): this demonstrated the expected temporal trends of expression for these genes, and indicated that those cells with the highest levels of *Gfra1* and *Id4* represent a basal state of development (**Figure 2B and S2**). We found that, in this most basal population, *Gfra1* and *Id4* expression are highly correlated (r=0.92), indicating that the regulation of both genes is highly synchronized, as opposed to a more complex structure of expression. Compared to *Gfra1*, we found that only 17 genes in the transcriptome show a more dramatic drop in expression as SSCs transition to a more restricted fate, including 5 transcription factors: *Utf1*, *Glis3*, *Fli1*, *Batf*, and *Foxf1* (**Figure S2I**).

Meiotic sex chromosome inactivation (MSCI) is an evolutionarily conserved phenomenon essential for proper spermatogenesis in mammals. During MSCI, transcription of sex chromosomes is silenced as part of a broader mechanism silencing unsynapsed chromatin (MSUC) ^13,14^. Consistent with previous reports we find that MSCI occurs at the start of pachytene (**Figure 2C**). Previous bulk RNA-seq studies suggested that some genes escape MSCI ^15,16^. However, our data do not support this conclusion, and instead indicate that the “escapees” are actually expressed after MSCI; thus it seems likely that the original finding was an artefact resulting from contamination of purified sub stages with later cells (**Figure 2D**).

Post meiotic cells have haploid genomes, meaning they have either an X or a Y chromosome but not both. However, cytokinesis does not fully complete in spermatogenesis resulting in synchronised chains of hundreds of cells, connected by μm-wide cytoplasmic bridges through which mRNA (or perhaps even mitochondria) could be shared ^17^. The extent to which mRNA sharing occurs is unknown, but it is a property of fundamental interest to evolutionary biology as most models predict a strong fitness benefit to fathers who can mask haploid selection in their gametes ^18^. Here, we find that, with respect to sex chromosome transcription, the genetically haploid cells are phenotypically diploid (**Figure 2E**) suggesting that cytoplasmic mRNA is efficiently shared, consistent with studies of individual genes ^19^. However, there remains a possibility that some genes are not shared, such as has been observed for the t-complex responder (Tcr) which functions as an antidote in the poison-antidote meiotic drive system of the t-complex ^20^.

Somatic cells in testis provide essential support to the germ cells throughout spermatogenesis but their precise functions have not been comprehensively delineated. Sertoli cells surround and nourish differentiating germ cells and phagocytose excess cytoplasm that is discarded by maturing spermatids. Leydig cells play a crucial role in germ cell maturation by maintaining testosterone production. Peritubular myoid cells, which are smooth-muscle-like cells, are involved in transporting immotile spermatozoa through the tubules. A number of studies suggested that somatic cells recognize different states of spermatogenesis and alter their functions accordingly, resulting in stage-specific patterns of transcription ^21^(Johnston et al.2008).

Additional subclustering on the 2 somatic cell clusters described above revealed heterogeneity within testicular somatic populations. We identified 11 different somatic sub-clusters — 5 Sertoli sub-populations, 3 Leydig sub-populations, 2 immune cell populations. (Macrophage and Lymphocyte) and 1 peritubular myoid cell population (**Figure S3**). All three Leydig sub-clusters (sub-clusters 6, 7 and 9) expressed genes *(Hsd17b3, Fabp3, Star, Insl3, Cyp11a1, and Cyp17a1*) involved in known Leydig-specific biological processes including testosterone biosynthesis from cholesterol, beta-oxidation of fatty acids, cholesterol biosynthesis, and retinoic acid metabolism. Sub-cluster 7 highly expressed genes involved in mitochondrial electron transport whereas cluster 9 was highly enriched with genes involved in detoxification and alcohol biosynthesis. All five Sertoli sub-clusters (sub-cluster 2, 5, 8, 10, and 11) expressed known Sertoli cell-specific markers (*Amhr2, Aard, Defb36, Cst12*). Based on the published work on stage-specific gene expression in Sertoli cells (Johnston et al.2008), four of these subclusters appear to represent different spermatogenic or seminiferous tubule stages. The gene expression pattern of sub-cluster 5 resembled that of seminiferous tubule stage V-VII, whereas sub-clusters 8, 10, and 11 resembled spermatogenic stage VII-XI, VIII-X and IX-XII, respectively (**Figure S3D**).

### Identification of gene modules using SDA

One specific challenge of analysing a developmental system is that cluster-based cell type classification might artificially define, hard thresholds in a more continuous process. Furthermore, a single cell's transcriptome is a mixture of multiple transcriptional programmes, some of which may be shared among cell types. In order to identify transcriptional programmes themselves rather than cell types, without *a priori* cell type classification, we applied sparse decomposition of arrays (SDA) ^22^. This is a model-based factor analysis method to decompose a matrix into sparse, latent factors, or “components” which identify co-varying sets of genes which, for example, could arise due to transcription factor binding or batch effects (**Methods**). When applied to single cell RNA-seq data, SDA generates two vectors of scores for each component: one reflecting which genes are active in that component, and the other reflecting the relative activity of the component in each cell, which can vary continuously across cells, negating the need for clustering (**Figure 3A-D**). This framework provides a unified approach to simultaneously soft cluster cells, identify co-expressed marker genes, and impute noisy gene expression (**Figure 3E, Figures S4 and S5, Methods**).

**Figure 3.**
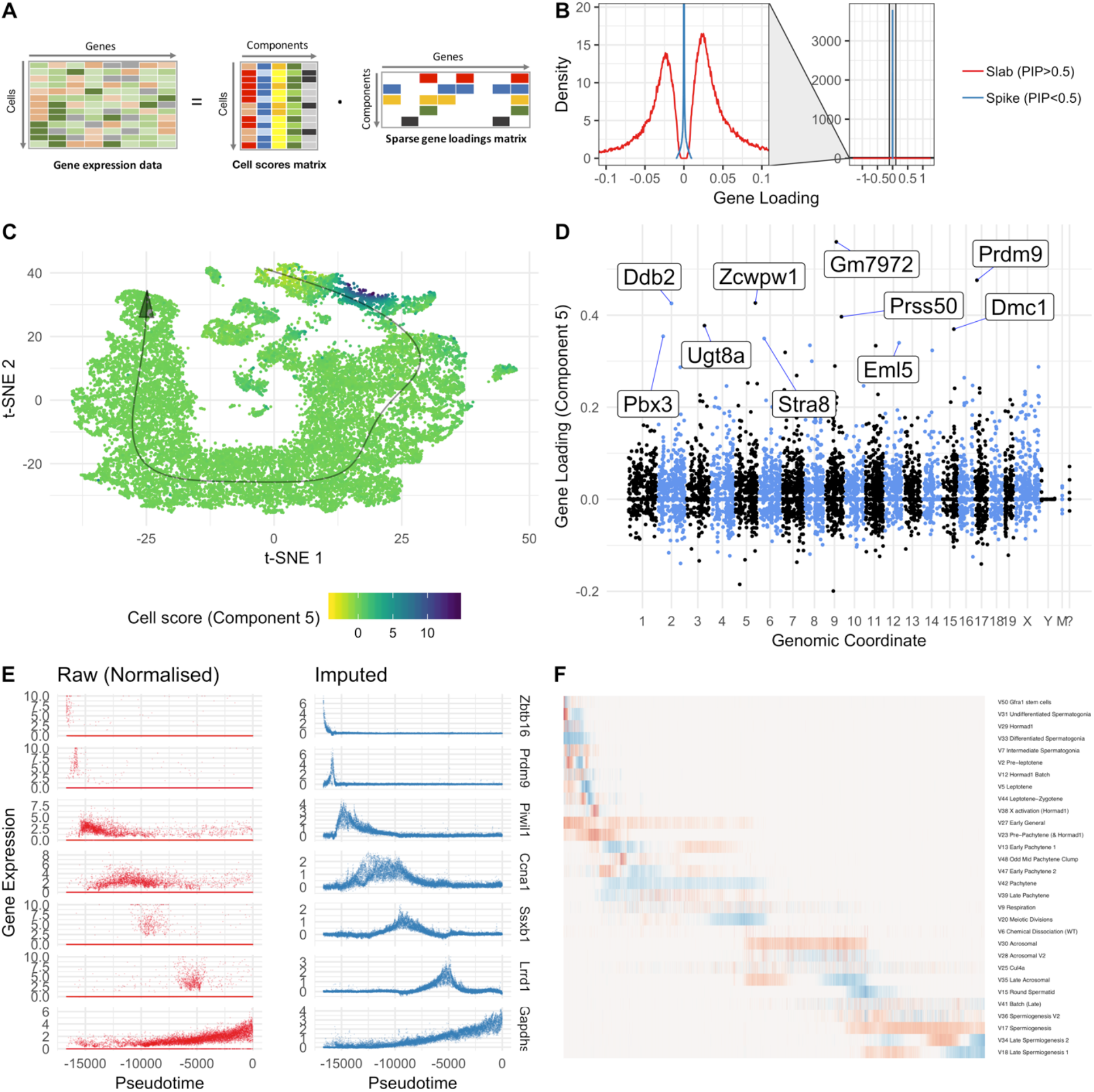
SDA identifies gene modules and maps them to cells. (A) We applied sparse decomposition analysis (SDA) to identify latent factors (“components”) representing gene modules. These components are defined by two vectors – one that indicates the loading of each cell on the component, and one that indicates the loading of each gene on the component. (B), SDA induces sparsity, so there are only few non-zero gene loadings on each component. (C), We identified 50 components from analysis of 20322 wildtype and KO cells (see also **Figure S4**). We illustrate interpretation of SDA components by visualization of component 5, a module of genes marking leptotene/zygotene. The loadings of component 5 in t-SNE space highlights a small set of cells at the expected early developmental stage of meiosis. (D), The component 5 loading of all genes in the transcriptome. Each point represents a gene loading. Genes are ordered by genomic location. Most genes have loading of 0, but a small number of genes have non-zero loadings including the well-known histone methyltransferase *Prdm9*. For detailed analysis of other representative components see **Figure S6**. (E), The model fit by SDA can be used to impute gene expression values for a given cell, conditional on that cell's estimated component loadings. Here we illustrate ability of imputation to improve the signal/noise ratio of expression for 7 genes with strong developmental regulation. We validated the utility of SDA-based imputation (Methods, **Figure S5**) (F), Cell component loadings for 30 germ components sorted by cell pseudotime. Each column corresponds to an individual cell and each row is a component. The component loadings are represented as a heatmap, with red colors representing positive loadings and blue representing negative loadings. Some components display both positive and negative loadings at distinct points in pseudotime. Although pseudotime information is not used in SDA analysis, SDA component loadings show coherent ordering throughout pseudotime.

We inferred 50 components using SDA: these represent all the major different cell types and developmental stages of spermatogenesis as well as batch effects and general processes such as respiration (**Methods**, **Figure S6**, **File S1**). Encouragingly, most components contained relatively few highly expressed genes (**Figure 3**) and showed directional biases in gene weightings (**Figure 3D**) not present in the model used for inference, but consistent with expectations for identifying a group of co-activated (or a group of co-repressed) genes. Overall, of 50 components, 6 represent batch effects, 5 are components with only a single cell, 13 are observed only in somatic cell types, 23 only in germ cells, and 3 components load on both somatic and germ cells. Within somatic cell components, we observe components corresponding to Sertoli cells (n=5), Leydig cells (4), macrophages (1), lymphocytes (1), peritubular myoid cells (1) as well as a component that seems expressed in all interstitial cells (1). Among germ cells-specific components, we observe components corresponding to processes active in spermatogonia (5), preleptotene spermatocytes (1), leptotene/zygotene (2), pachytene (5), diplotene (1), and spermiogenesis (7). Thus, we find multiple sub-components within existing recognised meiotic stages, adding considerable resolution relative to bulk-sequencing approaches. For some analyses below, we considered positively and negatively weighted genes within a component separately, in case these represent different modes of regulation, within the same groups of cells. After removing 5 batch effect components we were able to perform t-SNE and pseudotime construction with reduced technical noise resulting in improved resolution. Remarkably, although pseudotime information is not part of the SDA modeling framework, the cell loadings of SDA components show perfect agreement with the pseudotime assignment of each cell (**Figure 3F**).

One advantage of our approach is the ability to impute expression values, despite the sparsity of our single-cell data (mean 1,032 transcripts per cell), by combining data across cells, while still allowing individual cells to have their own scores. In individual cells, a lack of reads for many genes hinders our ability to estimate their expression. By performing imputation (**Figure 3E**) we are able to estimate expression of individual genes essential for meiosis even where zero reads are observed in a cell; via cross-validation, we verified that this approach uniformly improves our ability to rank genes according to their true expression levels, relative to using the raw read data (**Figure S5**).

We provide a web application to enable interactive exploration of gene expression and components at http://www.stats.ox.ac.uk/~wells/testisAtlas.html

### Germ Cell Components

Five components correspond to processes in spermatogonia. Markers of undifferentiated spermatogonia-*Zbtb16* (aka *Plzf*) ^8^, *Gfra1 ^10^*, *Nanos3 ^11^* and *Lin28a ^12^* – are spread through spermatogonial components 50 and 31, while component 33 contained genes involved in spermatogonial differentiation (**Table 1**).

**Table 1.**
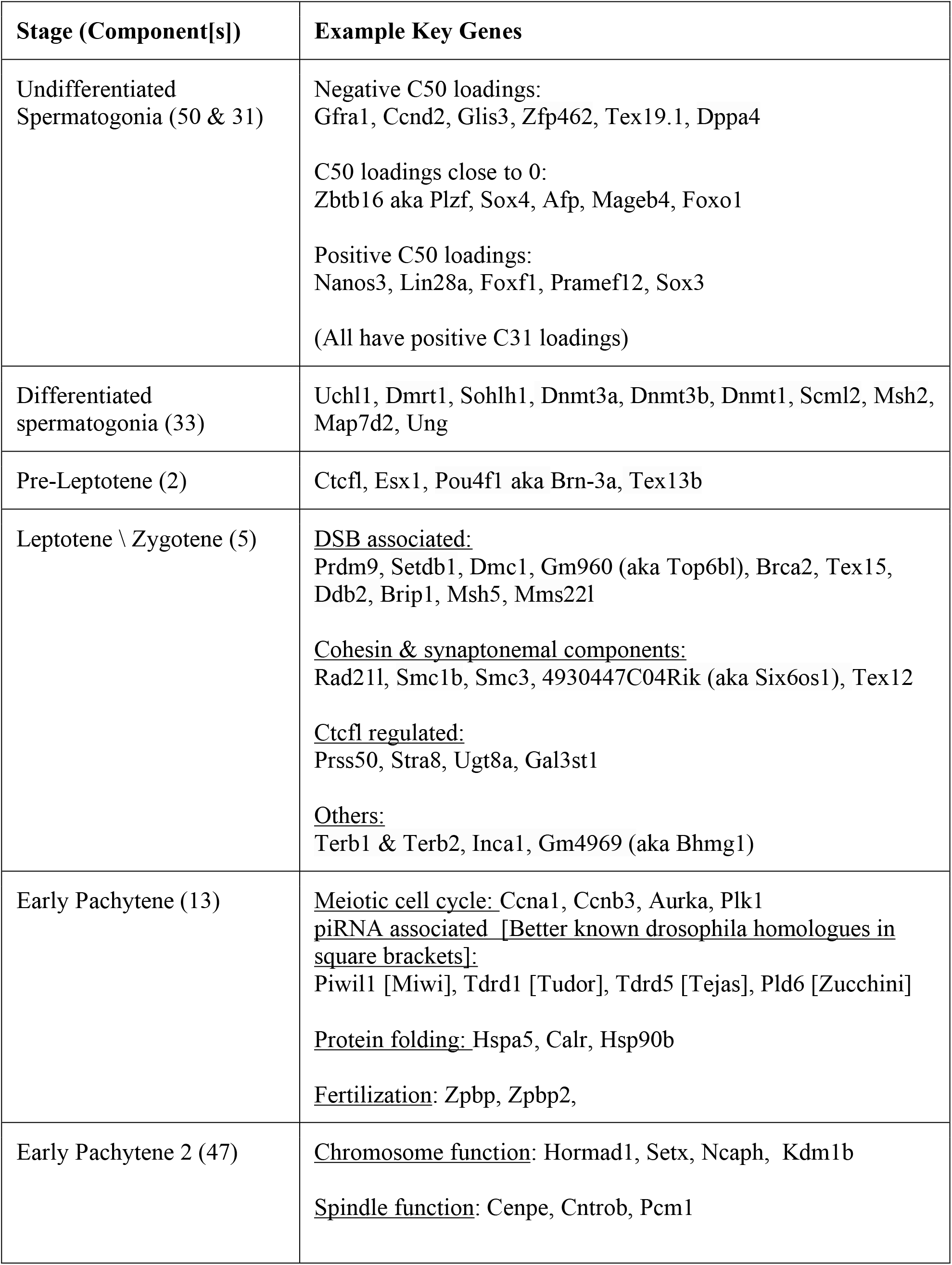
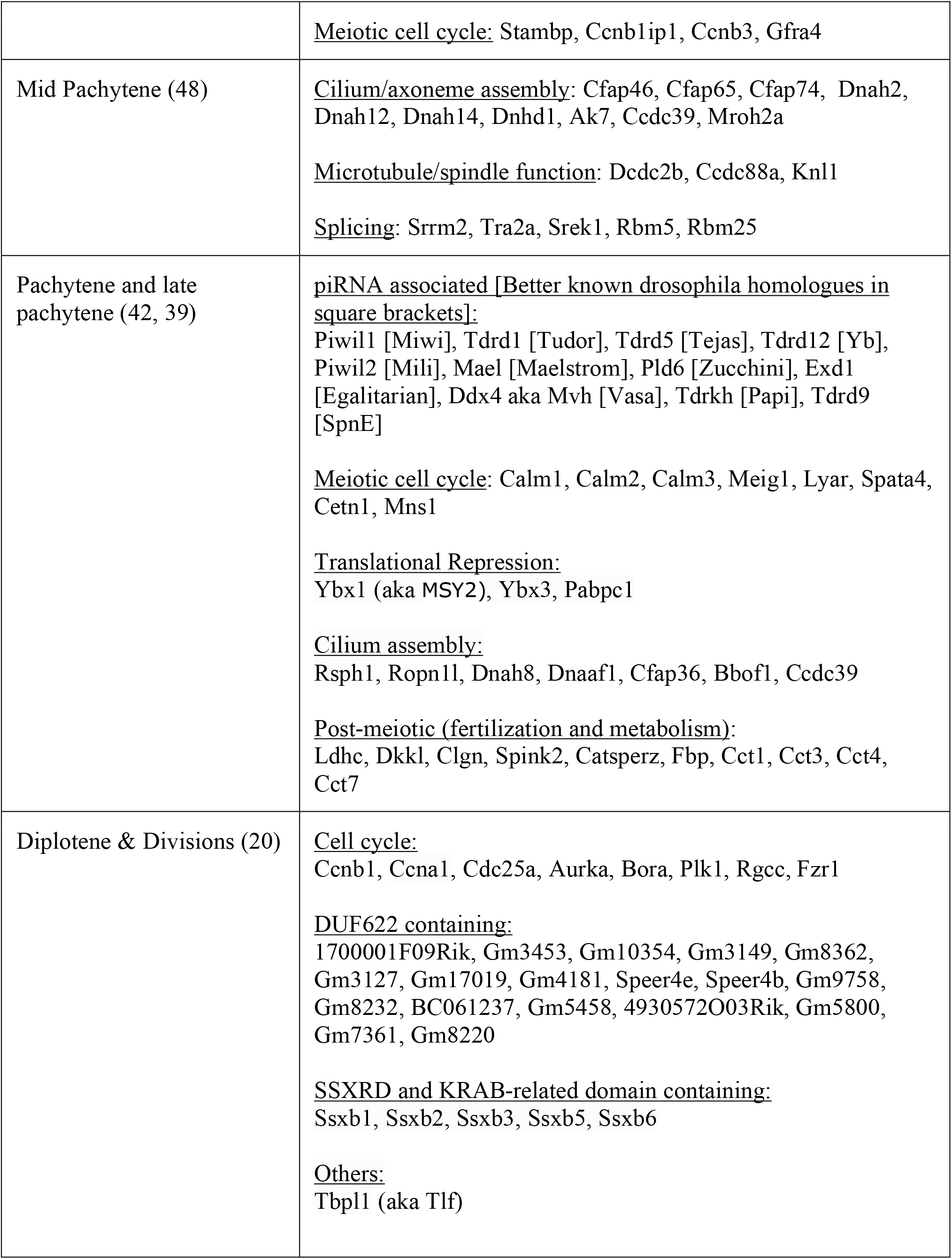
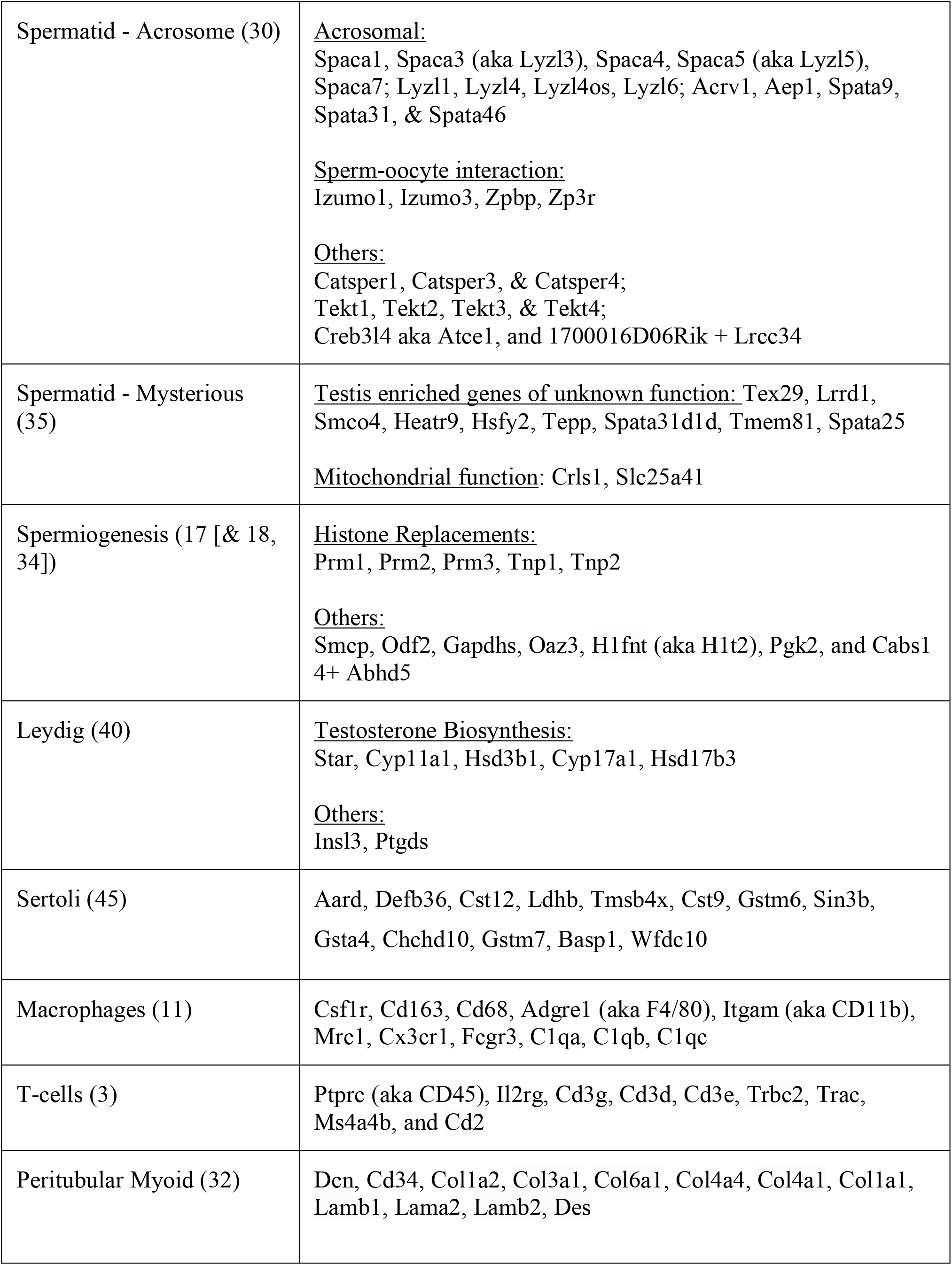
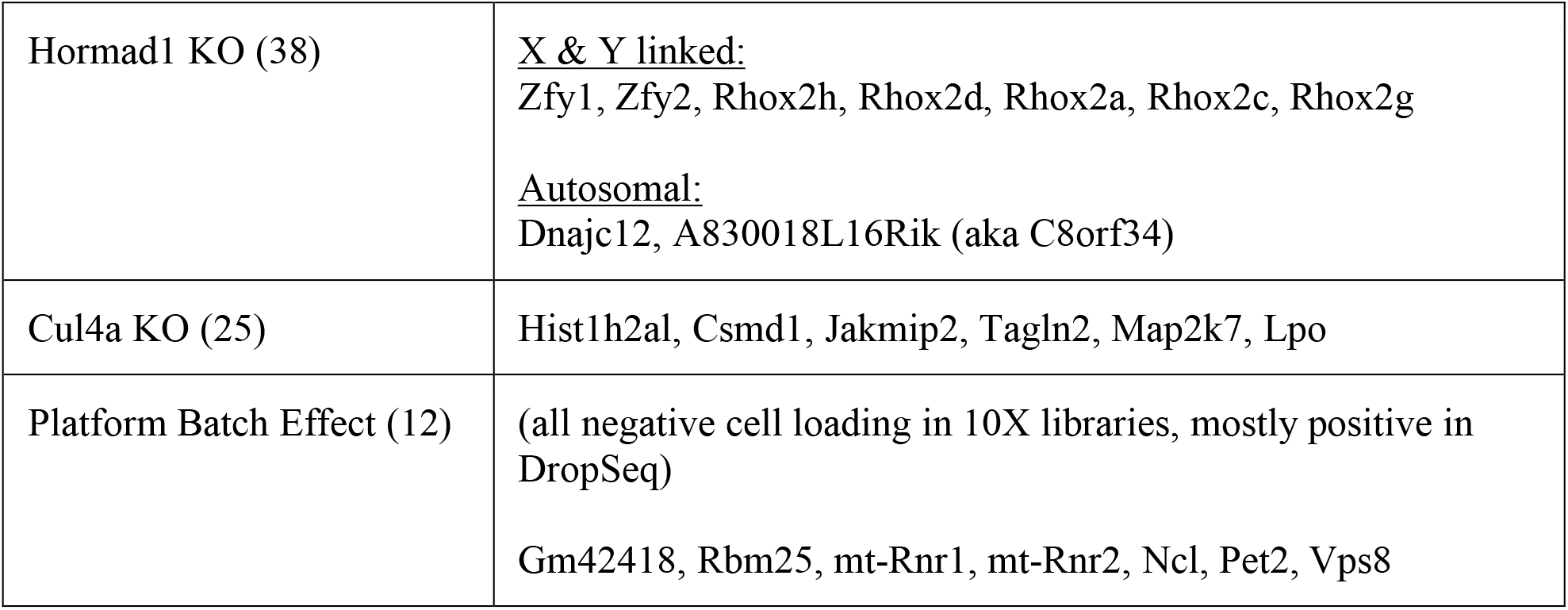
Key genes from example components of different stages

During meiosis there is an extended prophase I (lasting 14 days in mice), which is itself divided into a number of stages: Leptotene, Zygotene, Pachytene, and Diplotene. In Leptotene α Zygotene the homologous chromosomes undergo presynaptic pairing, aided by meiosis-specific cohesin, and induce several hundred programmed double-strand breaks at sites bound by *Prdm9*, the only known mammalian speciation gene ^23^; ^24^. We find a component with many of these genes, including *Prdm9* and components of the cohesin complex (Figure 3C-D; **Table 1**).

Previous approaches to germ cell transcriptional profiling have provided a single, static summary of pachytene expression from bulk sequencing of purified cells ^15,16^. Here, we are able to decompose pachytene gene regulation into 5 components (13, 39, 42, 47, and 48). These components have a striking lack of genes loading on the X or Y chromosome (**Figure S6E**), due to meiotic sex chromosome inactivation (MSCI). Although the cell loadings for these components overlap in pseudotime, they differ dramatically in their dynamics (**Figure 3F**). For instance, component 13 and 47 loadings appear to fluctuate, from positive, to negative, to positive again, while component 42 loading is constantly negative when active. Another interesting observation is that the genes with strong loadings within a component don't necessarily associate with a single, coherent functional process, nor even encode a set of transcripts that are all translated at the same point in spermatogenesis. Instead, components 13, 39, 42 and 48 each appear to involve both a substantial number of genes required for meiosis, as well as a second set of genes needed for some postmeiotic process, especially genes involved in sperm tail formation (**Table 1**).

The early pachytene components 13 and 47 are enriched for genes involved in the meiotic cell cycle (e. g.*Ccna1*, *Cdk1*), chromosome pairing and segregation (e. g.*Sycp3*, *Dmc1*, *Hormad1*), nuclear division (e. g.*Cenpe*, *Plk1*), and piRNA processing (e. g.*Tdrd1*, *Tdrd5, Tdrd9, Piwil1* and *Piwil2*). The next component in the sequence, 48, is an odd one restricted to a small part of t-SNE space and enriched for a large number of genes involved in axoneme/cilia assembly (many members of the *Cfap* family and dynein genes) as well as a smaller number of genes involved in microtubule/spindle formation (e. g.*Dcdc2b*, *Ccdc88a*, *Knl1*) and RNA splicing (e. g.*Srrm2*, *Tra2a*, *Srek1*). Component 42 and 39 (pachytene/late pachytene) are enriched for genes with similar biological functions - such as meiotic cell cycle, cilium assembly, piRNA processing, and translational suppression. These two components, as well as component 47, are significantly enriched for genes that are targets of the transcription factor MYBL1 (as determined by ChIP-Seq, **Figure S7B**).

Component 20 is particularly interesting. This component contains a number of genes known to be expressed in diplotene and or key regulators of cell division in addition to the *Ssx* family of genes (discussed further below) and also shows an enrichment of genes characterised by the presence of a DUF622 domain (18 in the top 88 genes) (**Table 1**). This fascinating gene family is rodent-specific and arose from duplication of the gene *Dlg5 ^25^*. It has previously been shown that many DUF622 genes experience similar epigenetic changes as the sex chromosomes during spermatogenesis, despite being located on the autosomes ^26^. Genes in this component are likely to be functional during meiotic divisions and perhaps the very first events thereafter.

We identified 7 post-meiotic components characterizing wildtype biology. Round spermatid component 30 contains many genes associated with the acrosome, an organelle which forms a nuclear cap containing hydrolytic enzymes used in fertilization ^27^ (**Table 1**). In addition, the gene *Lrrc34* has a high loading. We verified by immunofluorescence that the protein is indeed localised to the acrosome of round spermatids (**Figure 2A**).

Component 35, which is essentially concurrent to component 30 in pseudotime, is the most mysterious of all components that we detected. Dozens of protein-coding genes in this component are highly enriched in testis expression but have no known function (**Table 1**). This component also harbors a substantial number of genes with no apparent ortholog in humans. The existence of such a set of poorly characterized genes likely reflects the difficulty of studying postmeiotic male germ cells - these cells cannot be differentiated in vitro, contain numerous cell-type specific processes, and express many genes which are rapidly evolving.

The spermiogenesis components 17, 18 and 34 all contain many genes known to be expressed at the latest stages of spermiogenesis, before transcriptional arrest due to replacement of histones with protamines ^28^ (**Table 1**). In addition, *Abhd5* (aka CGI-58), a protein previously detected in testis lipid droplets ^29^, has high loadings specifically in these late components (17 & 18) and we show by immunofluorescence that it serves as an excellent marker of the residual body, a subcellular structure that has historically lacked a specific molecular marker. (**Figure 2A**).

### Somatic Components

In addition to components for the germ cell transcriptional programmes we identified components for at least 5 different somatic cell types (Sertoli, Leydig, Macrophages, T-cells, and peritubular myoid cells). Some components clearly mark multiple cell types that resolve separately in t-SNE space, while others mark groups of cells that may contain cryptic heterogeneity obscured by overlapping gene expression patterns. Leydig cells are responsible for the majority of testosterone production. Component 40 contains all of the key genes of testosterone biosynthesis within the top 55 genes, in addition to other well-known Leydig markers (**Table 1 and Figure S6G**).

Sertoli cells are responsible for nursing germ cells by providing nutrients and signalling, in addition to forming the blood-testis barrier and phagocytosing the residual body and apoptotic cells. Genes in component 45 have high correspondence (27 out of the top 30 markers) to previously discovered genes enriched in Sertoli cells (**Table 1**; ^30^.

Although the central section of the seminiferous tubules is immune privileged, in the interstitial space there are tissue resident macrophages (equivalent to Kupffer cells in liver or microglia in brain). Component 11 shows many key macrophage markers in the top 100 genes (**Table 1**). We also find a smaller group of T-cells represented by component 3 with high loadings for the T-cell markers (**Table 1**).

## *De novo* inference of transcription factor binding sites

The presence of many dynamic and finely tuned transcriptional programmes naturally leads to the question of their regulation. We used an existing approach^31^ to discover *de novo* motifs enriched in the promoter regions of the top 250 positive and negative genes (separately) for each component (**Methods**). We compared the resulting motifs with known motifs from the HOCOMOCO database, resulting in 16 groups of matched motifs, as well as one group of identified *de novo* motifs not clearly associated to a known motif, but most similar to the binding target of ATF1 (**Figure 4A, Figure S7A**).

**Figure 4.**
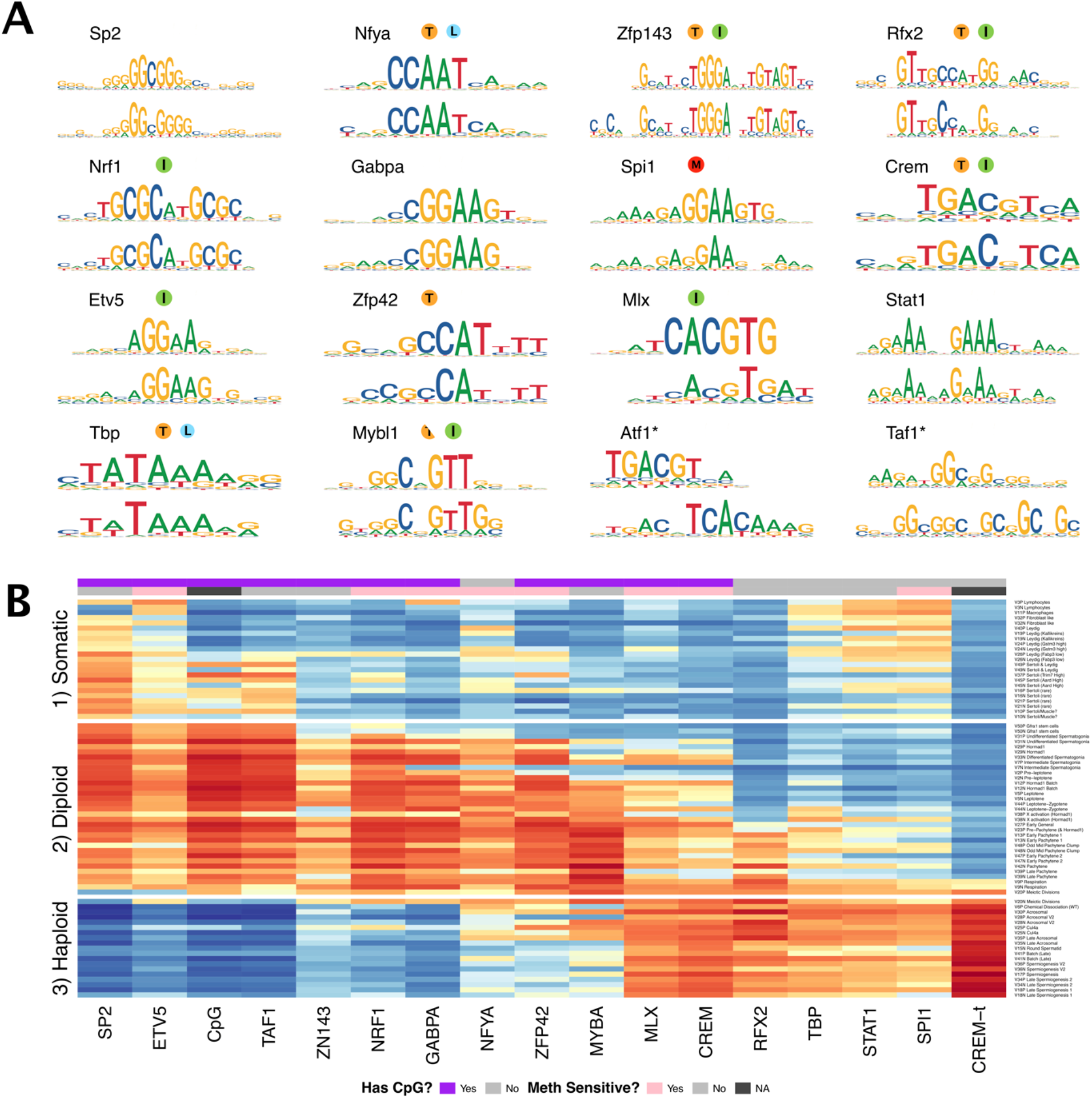
Components show shared *cis*-regulatory features. (A) Motifs discovered from the promoter sequences of genes with high component loadings. In each motif logo pair the lower logo shows the *de novo* inferred motif and the upper logo shows the motif in the HOCOMOCO database with the best match to the *de novo* motif. Orange “T” indicates this transcription factor is highest expressed in testis in GTEx database (half T indicates second highest). Green “I” indicates that a knockout of this gene produces an infertility phenotype. Blue “L” indicates a knockout of this gene is embryonic lethal. Red “M” indicates this gene is required for macrophage development.* after Atf1 indicates that we believe that the true transcription factor recognizing the *de novo* motif is not ATF1, but the tau isoform of CREM. (B) Association of gene loadings with the probability each *de novo* identified motif is found in the genes for each component. Coloring is an asinh scaled z-score from a correlation test between gene loadings and motif probabilities (calculated using the MotifFinder package from the denovo motifs. See also **Figure S7**), where red (blue) indicates positive (negative) association. The components (rows) are ordered by pseudotime. The column “CREM-t” shows the association with the motif probabilities using the *denovo* motif best matching Atf1 (i. e. the lower motif in the Atf1 pair in panel A). The additional column “CpG” shows association with number of promoter CpG dinucleotides for each component. Across the top of the panel, color bars indicate whether each motif contains a CpG, and whether the corresponding transcription factor is known to be methylation sensitive.

These include multiple well-known master regulators of spermatogenesis *Mybl1* ^32^, *Rfx2* ^33^, and *Crem* ^34^. In addition, we found *Spi1* (aka *PU.1*), a known master regulator of macrophage differentiation ^35^, specifically in the macrophage component 11, and distinct from the similar Stat1 motif that we observed in some later meiosis components. Analysis of ChIP-seq data for MYBL1, RFX2 and CREM validated our conclusion that covariation of expression in many components reflects a transcriptional programme (**Figure S7B**).

In our initial analysis we frequently identified the same motif (e. g. Sp2) in multiple components. Therefore, we looked at the association of motif presence/absence for particular genes with the component loadings, across all components for each motif (**Figure 4B**). This revealed an obvious “switch” with one group of transcription factors appearing to regulate early meiosis (prior to the meiotic division), and another group regulating meiosis post-division, with only the similar MLX and CREM motifs strongly spanning this divide. Moreover, most meiotic motifs spanned several components. This implies that promoter motifs might offer “broad-scale” control, but differences at “fine scales” among individual components might instead be driven by TF binding to more distant enhancer regions, mRNA degradation by microRNA, or other posttranslational mechanisms. Hence, additional work will be required to fully delineate the mechanisms controlling meiotic transcription.

Strikingly, all but one of the pre-division motifs as well as MLX/CREM contain CpG dinucleotides (**Figure 4**), with most, including the non-CpG exception (NFYA) also being sensitive to DNA-methylation in their binding^36,37^. None of the post-division motifs contain CpGs (p=0.007 by Fisher's exact Test for CpG occurrence). Indeed, we found an even stronger pattern of association simply using the count of CpG occurrences as a pseudo-motif (**Figure 4B**), indicating a major shift away from expression of genes whose promoters are CpG-rich as meiosis progresses.

To find potential effectors of this switch we looked in the component active at the stage of meiotic division, component 20. We found an enriched family of testis-specific proteins specifically expressed at this time characterized by the presence of both SSXRD and KRAB-related domains. The SSXRD domain has been studied in the context of synovial sarcomas where it was found to associate with the CpG binding protein CXXC2 (KDM2B) which is a component of a non-canonical polycomb complex ^38^. Interestingly another non-canonical polycomb component *Dcaf7* also has a high loading in component 20 ^39^. It has previously been observed that H3K27me3, a mark deposited by polycomb complex 2, increases dramatically between pachytene and the round spermatid stage ^40^. The KRAB-related domain has been studied as part of PRDM9 (the only other gene outside of the X-chromosome cluster to contain both the KRAB-related and the SSXRD domains), where it has been shown to interact with a number of proteins including the CpG binding CXXC1 ^41^, ^42^.

As we inferred these motifs *de novo* we were able to discover previously unknown motifs. Indeed, we identified a motif with similarity to the ATF1/CREM motifs in the late components, but with the addition of a CAA tail and lacking the central G nucleotide which would otherwise form a CpG dinucleotide (**Figure 4A**). We hypothesise that, plausibly, this may represent the binding motif of the tau isoform of CREM known to be active in late spermatogenesis ^43^. Consistent with the more general pattern of CpG occurrence we find this ATF1/CREM-t motif highly associated with post-division components; in fact it is the most strongly associated motif in multiple such components (**Figure 4B**).

## Characterization of testicular defects

The flexibility of the SDA modeling framework allows the identification of sets of genes that show significant covariation in extremely small numbers of cells. We reasoned that a joint analysis of mutant and wildtype cells using SDA would allow us to decompose expression variation into batch effects, normal gene regulation, and mutant-specific processes that reflect pathology. We selected four mutants to profile as models of male infertility – three mutants with known molecular mechanisms (knockouts of *Mlh3*, *Hormad1*, and *Cul4a*) as well as one knockin of a transgene (*Cnp*) that led to idiopathic infertility. Clear differences in histopathology were observed between testicular sections of wild-type and mutants (**Figure 5A**).

**Figure 5.**
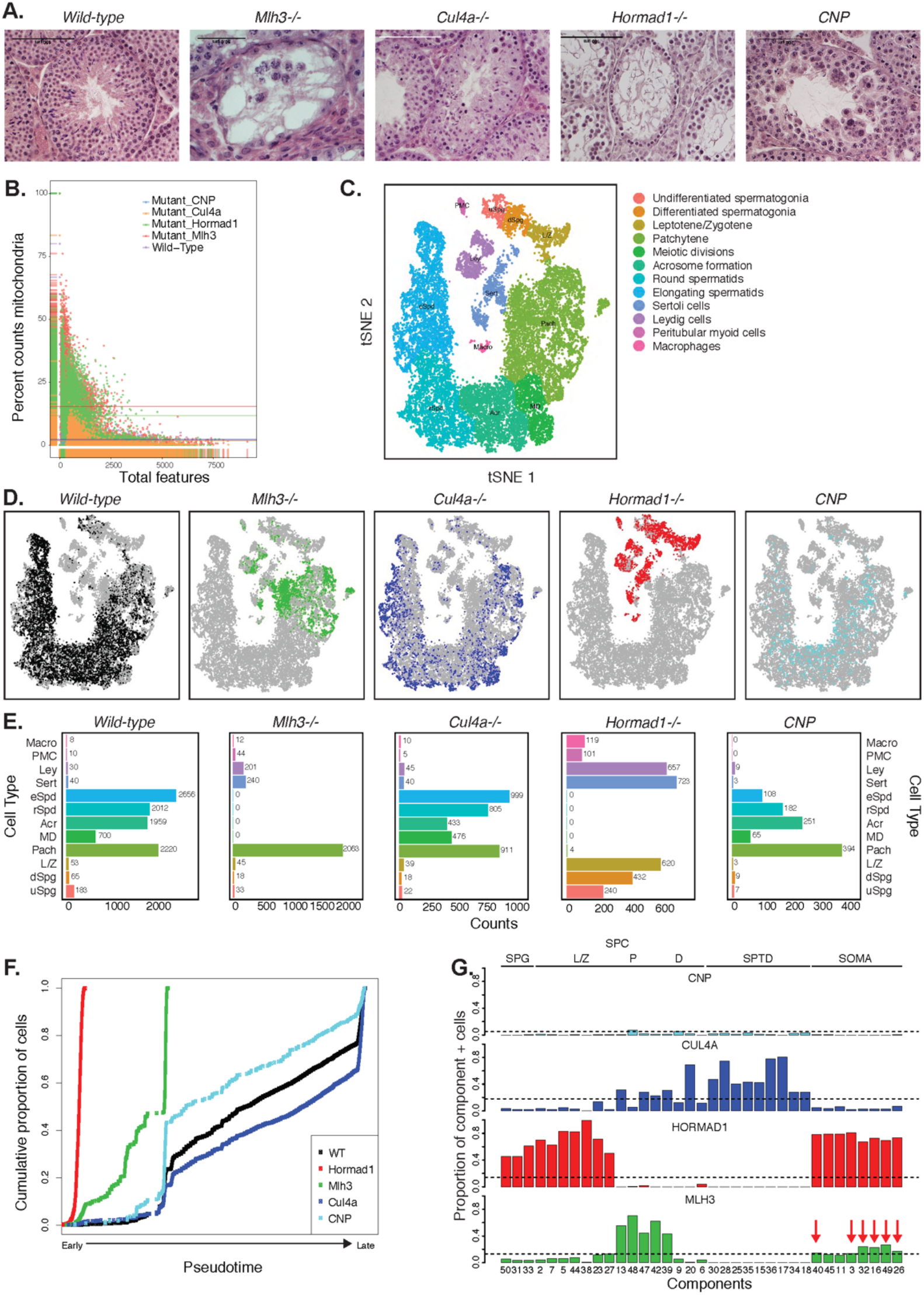
Characterization of mouse mutants with testicular phenotypes. (A) Histology sections from wildtype and mutant testis, illustrating the phenotypes observed in wildtype and the 4 mutant strains characterized by Drop-seq. Three of the strains, *Mlh3-/*-(REF), *Hormad1-/-* (REF) and *Cul4-/-* (REF) have known pathology, while strain *CNP* represents an unpublished transgenic line with spontaneous male infertility. (B) When compared to wildtype, cells from mutant strains exhibit significantly higher fractions of transcripts derived from mitochondrial genes, indicative of elevated rates of apoptosis. (C) We performed t-SNE clustering of both wildtype and mutant cells. All of the clusters observed by t-SNE on wildtype alone, were observed, but a few new clusters emerged from inclusion of mutant strains. See also **Figures S3** and **S4**. (D), Mutant strains occupied distinct locations within t-SNE space, reflecting both the absence of certain cell types in some strains (e. g.*Mlh3-/-* and *Hormad1-/-)*, as well as derangement of expression in remaining cells (e. g.*Hormad1-/-*). (E) Counting individual cell types provides a quantitative phenotype of cellular heterogeneity in each strain. (F) Cumulative distribution of cells along pseudotime from each mouse strain. The data clearly indicate that *Hormad1-/-* cells arrest prior to *Mlh3-/-* cells in the pachytene stage of spermatogenesis, while *Cul4a-/-* and *CNP* mice show quantitative deviation from WT in the abundance of postmeiotic cells. (G) As a way to summarize the SDA analysis of each strain, we plot the proportion of cells with strong component loadings from each strain separately. If cells are randomly distributed across components then we would expect the fraction of cells from each mutant to be the proportion of total cells sequenced from that mutant (dashed horizontal lines). Instead there are clear enrichments of component loadings in particular mutants, providing a fingerprint of pathology for those strains. Arrows indicate six components (3, 16, 26, 32, 40, and 49) that were enriched for cells from both *Mlh3-/-* and *Hormad1-/-* strains, but no other strains. SDA components are sorted by developmental stage, as indicated by horizontal lines across the top of the panel. SPG = spermatogonial components; L/Z = leptotene/zygotene components; P = pachytene components; D = diplotene components; SPTD = components in spermiogenesis; SOMA = somatic cell components.

Seminiferous tubules in *Mlh3-/-* and *Hormad1-/-* mice exhibited a complete early meiotic arrest and absence of spermatozoa.*Cul4a-/-* sections showed a partial impairment of spermatogenesis, indicated by a significant decrease in number of post-meiotic cells and abnormal spermatids. Sections from both *Cul4a-/-* and *Cnp* mice presented giant multinucleated cells but this type of cell was more prevalent in *Cnp* seminiferous tubules. Histological sections of *Cnp* mice displayed a clear defect in spermatogenesis as abnormal spermatids were observed; however, it is inconclusive which stage of spermatogenesis was affected without further molecular analysis.

After performing histological confirmation of testis defects in each animal to be sequenced, we generated 36,400 single cell profiles from mutant strains (**Table S1**). Cells from *Mlh3*-/- and *Hormad1*-/- animals showed higher rates of apoptosis compared to wild-type, *Cul4a* and *Cnp* (2% vs 14.5%, **Figure 5B**). We performed joint clustering and t-SNE visualization of all high-quality wildtype and mutant cells, which produced 32 distinct clusters. Based on differentially expressed markers in each cluster, we identified all subtypes of germ cells and somatic cell populations identified from the wildtype analysis, as well as 2 additional germ cell subtypes and 2 additional somatic cell types: undifferentiated spermatogonia, sex chromosome activated meiotic cells, immune cells, and peritubular myoid cells (**Figure 5C** and **Figure S2**).

Careful examination and quantification of cell-type composition differences in each mutant strain recapitulated the known pathology of mutants (*Mlh3-/-*, *Hormad1-/-* and *Cul4a-/-*) at an exquisite resolution. The location of mutant cells in t-SNE space illustrated absence of certain cell types within spermatogenesis. Consistent with the known biology, we observed that both *Mlh3-/-* and *Hormad1-/-* cells arrest at different stages of meiosis I; mid-pachytene and leptotene/zygotene respectively (**Figure 5D**). Derangement of certain cell types in the developmental trajectory was also observed as leptotene/zygotene *Hormad1-/-* cells and post-meiotic *Cul4a-/-* cells formed distinct clusters. By tallying counts of cells within each hard cluster, we generated a digital readout of the cellular composition of wildtype and mutant animals (**Figure 5E**).

Pseudotime analysis provided an even finer level of resolution for staging the time of onset of developmental problems in each strain (**Figure 5F**). By performing joint pseudotime analysis on all strains simultaneously, it will, in theory, be possible to fine map the timing of developmental defects. For instance, it is well known that it takes about 34.5 days for Type A spermatogonia to develop into mature spermatozoa ^44^. Given that our pseudotime-ordered set of 16,950 germ cells spans this entire developmental process, we estimate that the mean difference in developmental age between pseudotime-adjacent cells is 3 minutes. Although further work is needed to confirm a linear mapping of pseudotime to real time, we can use that mapping to estimate the difference in the mean time of arrest of *Hormad1-/-* cells and *Mlh3-/-* cells to be 12 days. This difference is reflected in the SDA components as well; *Mlh3-/-* animals have cells that load on pachytene components 47, 42 and 39, while *Hormad1-/-* animals do not.

HORMAD1 is a meiosis specific protein that regulates chromosome recombination, synapsis, and segregation. HORMAD1 normally marks un-synapsed chromosomes (such as the sex chromosomes). While HORMAD1 is removed by TRIP13 on synapsis, it persists on asynapsed chromosomes, which then undergo MSUC, leading to MSCI for the sex chromosomes ^45,46^. In *Hormad1-/-* spermatocytes, double-strand break formation and early recombination are disrupted as marked by the reduction of yH2AX, DMC1, and RAD51 foci. As expected, joint clustering analysis of wildtype and mutant cells indicated that all *Hormad1-/-* germ cells experience apoptosis at early pachytene of meiosis I due to checkpoint failures. Along with this arrest phenotype, the *Hormad1-/-* leptotene/zygotene cells form two distinct clusters outside of the leptotene/zygotene cells of all other strains (Clusters 30 and 32, **Figure S2**). A list of significant differentially expressed genes between the two clusters included a number of sex chromosome genes **(Table S2**). Consistent with these observations, we found one SDA component (38) with much higher loadings on the sex chromosomes than autosomes (**Figure 6A, Table 1**), and for which those cells showing significant loadings are all *Hormad1*^-/-^. We find that not only does *Hormad1^-^*^/-^ fail to silence previously expressed sex-linked genes, many previously *un*expressed sex-linked genes such as *Rhox2h* have high expression (**Figure 6B**). Interestingly, there are also multiple autosomal genes with high loadings. This may be due to ectopic expression of sex-linked transcription factors; for example, *Zfy1* and *Zfy2* were previously shown to cause pachytene arrest when misexpressed ^47^. We find a very strong association between genes in this component and genes overexpressed in mice which have mutations in either *Hormad1* (p<10^-35^) or *Trip13* (p<10^-150^) ^48^**; Figures S6A-D**).

**Figure 6.**
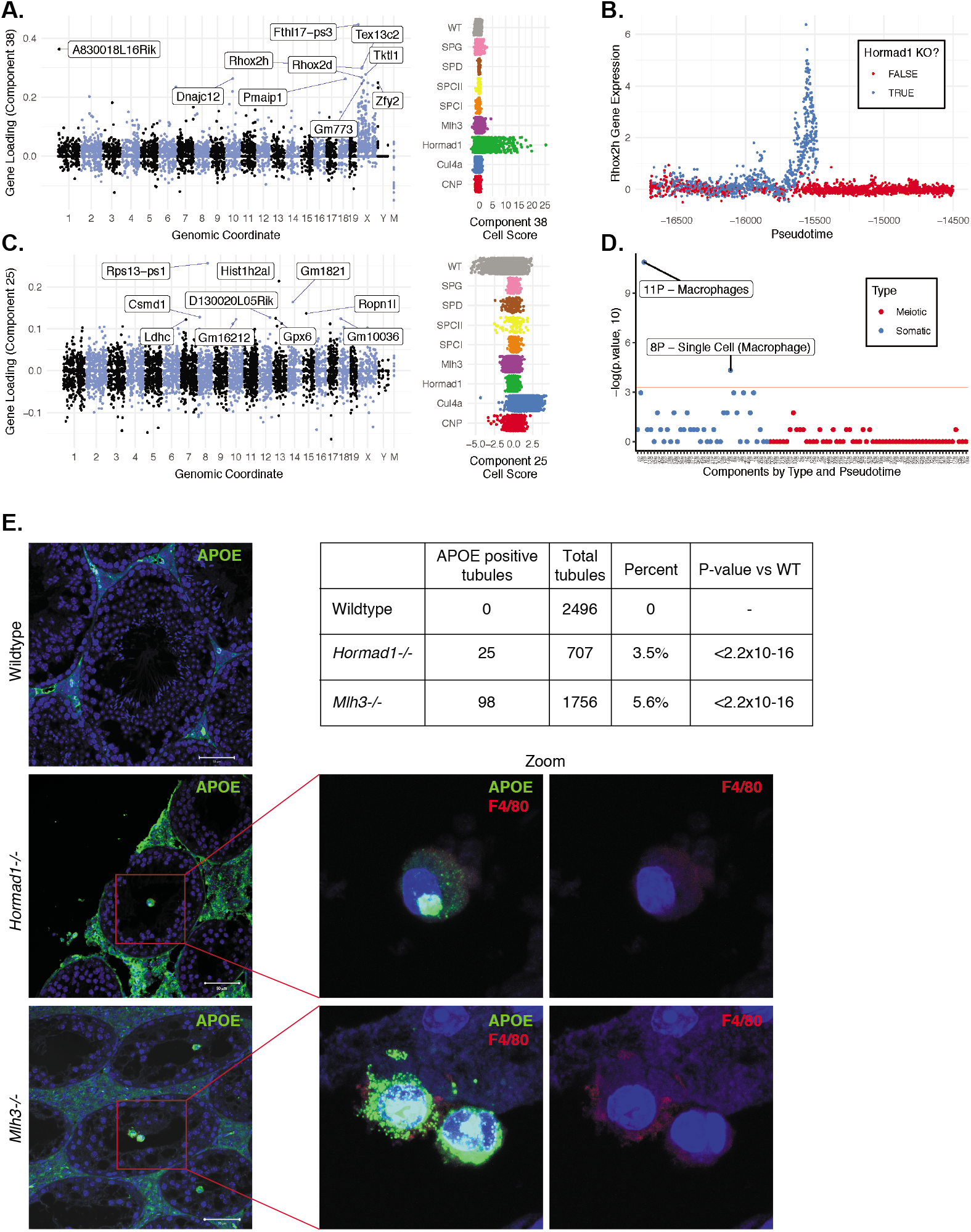
Dissection of strain-specific pathology. (A) SDA component 38 is comprised largely of genes on the X chromosome, with a gene loading direction that indicates failure of X inactivation. As illustrated by the cell scores (loadings) for this component, it is restricted to *Hormad1-/-* cells. (B) Pseudotime analysis indicates that *Hormad1-/-* cells diverge developmentally from all other strains around leptotene/zygotene. In this illustration, the X-linked gene *Rhox2h* is shown to have low or no expression in all cells prior to meiosis, and then rapidly increased expression specifically in *Hormad1-/-* cells until this lineage arrests. (C) Component 25 is the component most strongly enriched for *Cul4a-/-* cells. (D) We identified 6 components with shared enrichment for both Mlh3-/- and Hormad1-/- cells; these components contained genes with numerous significant GO associations related to Alzheimer's disease (AD) pathology (main text, **Figure 5G**). For each SDA component, we tested for association between known AD genes and genes with either positive (P) or negative (N) loadings on that component. AD genes are highly enriched for expression in component 11, corresponding to macrophages. (E) Further investigation of protein expression of AD genes revealed APOE+ (green) cells within the tubules of *Mlh3-/-* and *Hormad1-/-* but not WT. These cells showed nuclear morphology different from native germ cells or Sertoli cells, and stain positive for the macrophage marker F4/80. The inset table summarizes raw data on the frequency of APOE+ tubules obtained by microscopy. The frequency of APOE+ tubules is more common in each mutant strain when compared to WT by Fisher's exact test. Scale bar = 50mm.

MLH3 is an essential protein required for crossover formation in early meiosis and for binding of MLH1 to meiotic chromosomes. Studies on *Mlh3-/-* testes have shown depletion of spermatocytes and some spermatogonia due to apoptosis in diplonema induced by a reduction of chiasmata and a loss of recombination nodules ^49^. Interestingly, in contrast to *Hormad1-/-*, we found no obvious transcriptional phenotype in *Mlh3-/-* cells either by SDA analysis or by comparison of expression levels between hard-clustered wildtype and mutant cells (other than differential expression of *Mlh3*). Instead, *Mlh3-/-* spermatocytes might simply trigger apoptosis through existing checkpoint protein machinery assembled earlier in development. Using the simple pseudotime analysis described above, we can estimate that if a transcriptional response was triggered, it must last less than 33 minutes for it to be missed in our sample of cells (**Figure 5F**).

CUL4A is a major component of the E3 ubiquitin ligase complex called CRL4 which is known to regulate cell cycle, DNA replication, DNA repair, and chromatin remodelling ^50^. Studies on *Cul4a-/-* mice noted that some spermatocytes arrest at the pachytene stage of meiosis I induced by the pachytene checkpoint, whereas remaining spermatocytes complete meiosis but the resulting spermatozoa present oligoasthenospermia and severe malformations ^51^. The molecular basis of observed abnormal phenotypes in spermatozoa remains unclear. Our joint analysis revealed that pre-meiotic and meiotic *Cul4a-/-* cells clustered well with wildtype cells but certain phases of post-meiotic cells formed distinct clusters on the t-SNE plot (**Figure 5D**). We identified a single SDA component (25) that was highly specific to *Cul4a-/-* cells (**Figure 6C and Table 1**). This component corresponds to dozens of genes that are overexpressed in *Cul4a-/-* mutants when compared to all other strains, with GO enrichments related to spermatid development, motility and capacitation. These findings are consistent with the observed phenotype of *Cul4a-/-* mice and provide new leads to investigate mechanisms of pathology.

The cells from *Cnp* mice did not form distinct clusters, nor did they show SDA component loadings distinct from wildtype cells. Although the presence of multinucleated giant cells, hypocellular seminiferous tubules and infertile phenotype point to a serious defect in spermatogenesis, it is difficult to determine which stages are affected using single-cell expression data. One possible explanation of missing important biological signals may be that *Cnp* mice presents a partial arrest phenotype which masked the developmental abnormalities. Another possible explanation is that droplet-based sequencing library preparation may undersample the cells with aberrant transcriptional signatures.

## A convergent phenotype of meiotic arrest mutants

Despite the differences in cell composition or component loadings among mutant strains, we identified 6 somatic components -3, 16, 49, 40, 26, and 32-showing a specific enrichment for *Mlh3-/-* and *Hormad1-/-* cell loadings when compared to all other strains (**Figure 5G**). Three of these components, 16, 49 and 26, were highly enriched for genes involved in amyloid-beta formation, metabolism, and clearance, including *Apoe*, *App* and *Clu* (**Table S3**). Excessive production of amyloid-beta, which is a primary cause of Alzheimer's disease, was not previously reported in these mutants, and the possible physiological role of such production is unclear. We tested multiple antibodies to human amyloid-beta that failed to work on our tissue. To further evaluate the expression of Alzheimer's disease (AD)-related genes across all 5 mouse strains, we tested individual SDA components for enrichment of expression of AD risk genes identified in a recent GWAS, identifying component 11 (macrophages) as specifically and strongly enriched (p< 10^-12^, **Figure 6D** and **Figure S6F**). Immunofluorescence staining for the protein product of one well-studied AD gene, *Apoe*, in wildtype animals showed low levels of specific staining confined to the interstitial space (**Figure 6E**). Both *Mlh3-/-* and *Hormad1-/-* displayed interstitial cell with more intense staining of APOE, as well as a greater abundance of APOE+ cells. Most remarkably, we found a rare population of APOE+ cells within the tubules of *Mlh3-/-* and *Hormad1-/-* that are never observed in wildtype. We screened 4,959 tubule cross-sections to establish more precise estimates of APOE+ cell frequency in these three lines (**Methods**). When compared to the frequency in wildtype tubules (0/2496 tubules), we see higher frequencies of intratubular APOE+ cells in *Mlh3-/-* (25/707 tubules, 3.5%, p< 2.2 x 10^-16^) and *Hormad1-/-* (98/1756 tubules, 5.6%, p< 2.2 x 10^-16^). These APOE+ cells displayed a nuclear staining and morphology that are distinct from normal germ cells and Sertoli cells, and appeared more similar to APOE+ cells outside of the tubules. These cells stained for F4/80, a well-established macrophage antigen, which was quite surprising, as it is widely believed that only preleptotene spermatocytes transit the blood-testis barrier ^52^; ^53^. Co-staining of F4/80 with an antibody for activated CASPASE-3, a marker of apoptosis, failed to identify any double positive cells, so we exclude the possibility that intratubular F4/80 protein expression was somehow an artefact of an apoptotic cell population. A deep review of the literature indicated that intratubular macrophages have been described previously, always in the context of testicular disease ^54^; ^55^. The mechanisms by which macrophages transit the blood-testis barrier, and the corresponding cues for migration, await further investigation.

## Discussion

The extensive cellular heterogeneity of the testis has limited the application of genome technology to the study of its gene regulation and pathology. Here, we described how the SDA analysis framework can be applied to single-cell RNA-sequencing data of the testis to overcome the challenge of heterogeneity by summarizing gene expression variation into components that reflect technical artifacts, cell types, and physiological processes. Rather than clustering groups of *cells*, SDA identifies components comprising groups of *genes* that covary in expression, and represents a single cell as a sum of such components. This revealed previously uncharacterised complexity, with multiple different components even within recognised meiotic stages such as pachytene. We also identified components corresponding to highly specific and interpretable pathology in one or more mutant strains.

By performing *de novo* motif analysis, we observed that it is possible to identify transcription factors critical for the meiotic program without prior knowledge, as well as other motifs not currently well characterised. It seems likely that our analysis of promoters is only a first step towards what is possible here via – for example – analysis of enhancers and other regulatory sequences, and we hope that future data will allow this, working towards identifying the full set of transcription factors, and their targets, used in mammalian spermatogenesis. The apparent dramatic change away from the use of factors binding CpG dinucleotides, and whose binding is often disrupted by methylation of such dinucleotides, after the first meiotic division, is one area for such further research – whether this involves *Ssx* genes, DUF622-containing genes, or other factors. More generally, in combination with temporal information from pseudotime analysis, it will be possible to create a model of the cascade of gene regulation, and by comparison across species, better understand the constraints on the precise timing and ordering of regulatory events.

Finally, we note that gene expression components (for example those identified by SDA) represent an attractive way to build a dictionary of pathology of the testis. The construction of new component models using a larger panel of mutants with known pathology will accelerate the interpretation of idiopathic mutants, and, ultimately, could provide a framework for a much more advanced diagnosis of human infertility than is currently in practice.

## Acknowledgments

We thank Abul Usmani for assistance with animal husbandry and advice on *Pou5f1:GFP* mice, Jeffrey Milbrandt and the WashU Genetics Department Single Cell Program for support, Liang Ma for providing *Cul4a*-/- mice, Joe Dougherty for providing *Cnp* mice, and Katinka Vigh-Conrad for assistance with figures. We also thank the Alvin J. Siteman Cancer Center at Washington University School of Medicine and Barnes-Jewish Hospital in St. Louis, MO, for the use of the High-Speed Cell Sorter Core, which provided cell sorting service. The Siteman Cancer Center is supported in part by an NCI Cancer Center Support Grant #P30 CA91842. This work was supported by National Institutes of Health Grants R01HD078641 and R01MH101810 to D. F. C., and Wellcome Trust grants 098387/Z/12/Z to S. M. and 109109/Z/15/Z to D. W.

## Author Contributions

M. J. and J. R. performed all Drop-seq experiments; D. W., M. J., D. C. and S. M. analyzed the data. J. R., S. A. and M. J. performed histology. D. W., M. J., S. M. and D. C. wrote the paper with input from all authors. D. C., J. M. and S. M. supervised the project.

## MATERIALS AND METHODS

### Mice

All animal experiments were performed in compliance with the regulations of the Animal

Studies Committee at Washington University in St. Louis under protocol #20160089. Mice were housed in a barrier facility under standard housing conditions with *ad libitum* access to food and water and a 12hr:12hr light/dark cycle. All single-cell RNA sequencing experiments were carried out with sexually mature animals (ages of mice in this paper vary from 11-38 weeks) except for *Pou5f1-EGFP* transgenic animal testes which were collected at post-natal age (P) 7. For specific age of mouse at the time of testes collection for different batches, please refer to **Table S1**.

Samples for histological studies were also collected at the time of testes collection for single-cell RNA sequencing. The mouse lines used in this paper are the following:

1. C57BL/6J male mice were used for Hoechst-FACS and total testis single-cell RNA sequencing experiments.
2. B6;CBA-Tg(*Pou5f1*-EGFP)2Mnn/J reporter mice were used for enriching and isolating spermatogonia type A cells. Testes from five mice at post-natal age P7 were pooled to generate single-cell suspension and FACS sorted for GFP positive cells, followed by Drop-seq.
3. B6.129-*Mlh3^tm1Lpkn^*/J heterozygotes were bred to maintain the colony and male homozygotes were used for Drop-seq experiments.
4. B6;129S7-*Hormad1^tm1Rajk^*/Mmjax heterozygotes were bred to maintain the colony and male heterozygotes were used to Drop-seq and 10X Chromium experiments.
5. B6;129 *Cul4a*^-/-^ mice were used for generating Drop-seq data
6. C57BL/6J CNP-EGFP BAC-TRAP mice were used for Drop-seq data

### Single-cell Suspension Preparation

#### Mechanical Dissociation of Testes

Two different types of testicular dissociation protocols were used in this paper: enzymatic and mechanical. Both enzymatic and mechanical protocols were previously published in ^56^ and ^57^. These methods were modified appropriately for single-cell RNA sequencing.

For mechanical dissociation method, fresh testes were decapsulated in 1X DMEM and cut into small pieces (approximately 2-3mm^3^). These tissue fragments were transferred to a 50μm Medicon, a tissue disaggregator and tissue fragments were dissociated in 1mL 1X DMEM for 5 minutes on Medimachine. The resulting single-cell suspension was aspirated from Medicon with a 3mL needless syringe. This dissociation/aspiration step was repeated three times and total of 3mL single-cell solution was retrieved. Then the single cells were filtered through sterile 40um strainers twice and triturated for 1 minute with a wide orifice disposable Pasteur pipet. Cells were spun down at 500xg for 10 minutes at 4°C and re-suspended in 2mL 1X DMEM. Finally, cells were filtered once more with sterile 50um filter, adjusted to 100 cells/μl concentration, and placed on ice until processed for Drop-seq or submission to Genome Technology Access Center at Washington University in St. Louis for 10X Genomics Chromium service. Single-cell RNA sequencing experiments were performed within ~30 minutes of testes collection for mechanical dissociation.

#### Enzymatic Dissociation of Testes

Solutions necessary for enzymatic dissociation were prepared fresh prior to testes collection and these solutions are as follows: 120U/mL collagenase type I in 1X DMEM; 50mg/mL trypsin in 1mM HCl; 1mg/mL DNase I in 50% glycerol. For enzymatic dissociation method, decapsulated fresh testes were collected in 15mL conical tubes, one testis per tube. Each testis was dissociated in 6 mL of collagenase type I solution and 10μl of DNAse I solution with horizontal agitation at 120rpm for 15 minutes at 37°C. Tubules were decanted for 1 minute vertically at room temperature and supernatant was discarded. Another 4mL of collagenase type I solution, 50μl of trypsin solution and 10μl of DNAse I solution were added to each tube and incubated with horizontal agitation at 120rpm for 15 minutes at 37°C. Testicular tubules were triturated with a plastic disposable Pasteur pipet with a wide orifice for 3 minutes. Another 30μl of Trypsin solution and 150μl of DNAse I solution were added and incubated for 10 minutes with horizontal agitation at 120rpm. Then 400μl Fetal Bovine Serum (FBS) was added to deactivate dissociation enzymes. Finally, collected single-cell suspension was passed through 40μm filter twice and stored on ice until processing for Drop-seq. These cells were processed within 1.5 hour of the testes collection.

For digesting *Pou5f1*-EGFP mice testes, we adapted a protocol described previously ^58^. Briefly, testicular tubules/fragments were incubated in 200μg/mL trypsin solution for 15-20 minutes with intermittent pipetting followed by 300μl FBS addition for inactivating trypsin. Single-cells suspension was filtered through 50μm filters twice and stored on ice until FACS.

#### Isolation of Germ Cell Populations by Flow Cytometry

##### Hoechst-FACS for spermatocytes and spermatids

For isolation of major germ cell populations, we adapted a Hoechst-FACS protocol and sequential gating strategies described in (Lima et al.2017). Briefly, 10μl Hoechst and 2μl of propidium iodide (PI) were added to single-cell suspension obtained from one testis and incubated at room temperature for 20 minutes. Then single-cell suspension was filtered through a 50um cell strainer. Cells were sorted and analyzed using Beckman Coulter MoFlo Legacy cell sorter and Summit Cell sorting software. First, debris were excluded based on forward scatter (FSC) and side scatter (SSC) plot pattern. Single cells were gated by adjusting FSC and pulse width threshold. Dead cells were gated and removed based on PI intensity. A minimum of 500,000 events were observed before proceeding to gating on different germ cell populations. Then, cell count histogram was plotted based on Hoechst blue fluorescence and observed three peaks, representing haploid (1C), diploid(2C), and tetraploid (4C) populations. Then Hoechst-blue and Hoechst-red fluorescence intensities were plotted to refine spermatocytes and spermatids populations.

##### Spermatogonia type A

For isolating spermatogonia type A cells from the *Pou5f1-EGFP* reporter mice, cells were analyzed and sorted with the same cell sorter and software described above section. Similar sequential gating strategies were followed. Debris were excluded, single cells were gated and dead cells were excluded. Then, GFP+ cells were gated on a plot of GFP vs FSC.

##### Single-cell RNA Sequencing Library Generation

###### Drop-seq Procedure

Drop-seq sequencing libraries were generated according to a protocol described previously (Macosko et al.2015). Cells and beads were diluted to co-encapsulation occupancy of 0.05. Two bead lots were used for generating Drop-seq data (For more details, see **Table S1**). Individual droplets were broken by perfluorooctanol, followed by bead harvest and reverse transcription of hybridized mRNA. After Exonuclease I treatment, aliquots of 2000 beads were amplified for 14 PCR cycles (all necessary PCR reagents and conditions were identical to Macosko et al.2015). PCR products were purified using 0.6x AMPure XP beads and cDNA from each experiment was quantified by Tapestation analysis.600pg of cDNA was tagmented by Nextera XT with the custom primers, P5_TSO_Hybrid and Nextera 70X. The single-cell sequencing library from each batch was either pooled with another batch or sequenced separately on the Illumina HiSeq2500 at 1.4pM or MiSeq at 8pM, with custom priming (Read1CustSeqB Drop-seq primer).

###### 10X Genomics Procedure

A single-cell suspension from a *Hormad1-/-* mouse was processed through the 10X Genomics Chromium instrument according to the manufacturer's protocol. Briefly, single cells were added to microfluidics chip channel along with oil and barcoded gel beads and sorted into droplets. Within the gel beads emulsion, cells were lysed, RNAs were reversed transcribed, cDNA were amplified and sequencing libraries are constructed. Generated single-cell sequencing libraries were sequenced on Illumina HiSeq2500.

###### Collection and Processing of Testes

For histological studies, testes were collected in 4% paraformaldehyde (PFA), incubated overnight at 4°C and washed with 70% ethanol. For hematoxylin and eosin staining, testes were collected in modified Davidson fixative and after 24-hour incubation at room temperature, tissues were transferred to Bouin's solution for another 24-hour incubation at room temperature. Fixed testes were dehydrated through a series of graded ethanol baths and embedded in paraffin. Then 5μm sections were cut on clean glass slides.

###### Hematoxylin & Eosin (HE) Staining

Hematoxylin & Eosin staining was performed on each mouse line (Wildtype, *Mlh3-/-, Hormad1-/-, Cul4a-/-*, and *CNP-EGFP*) to assess overall morphology of testicular tissue. Slides were deparaffinized with xylene and rehydrated through a series of graded ethanol bath to PBS. Standard HE staining protocol was adapted from Belinda Dana (Department of Ophthalmology, Washington University in St. Louis) and followed with Hematoxylin 560 and 1% Alcoholic Eosin Y 515.

###### Immunofluorescence Staining

Prior to immunofluorescence staining, antigen retrieval was performed by boiling slides in citric acid buffer for 20 minutes and tissue sections were blocked in blocking solution (0.5% Triton X-100 + 2% goat serum in 1X PBS) for an hour at room temperature. Primary antibodies were diluted to antibody-specific dilution (see Key Resources Table) and incubated overnight at 4°C in a humid chamber. Then, slides were incubated in secondary antibodies (1:300 dilution) at room temperature for 4 hours in a humid chamber. After the secondary antibody incubation, sections were stained with Hoechst (1:500 dilution), washed with 1X PBS and mounted with ProLong Diamond Antifade Mountant for visualization under confocal microscope.

###### Preprocessing of Drop-seq Data

Paired-end sequencing reads were processed, filtered and aligned as described in Macosko et al.2015. The specific steps and tools for this process is further outlined in Drop-seq Computational Cookbook (http://mccarrolllab.com/wp-content/uploads/2016/03/DropseqAlignmentCookbookv1.2Jan2016.pdf). STAR aligner was used to map the processed reads to mouse genome. A STAR indexed genome was generated using mm10 mouse genome and GRCm38 gene annotation (release version 76) with default setting. Following the alignment, digital gene expression (DGE) matrices were generated for each experimental batch.

###### Preprocessing for 10X Chromium Data

One *Hormad1-/-* mutant data was generated from 10X Genomics Chromium. Cellranger toolkit was used to de-multiplex, align reads to mm10 genome, collapse UMIs and generate gene-cell matrices. This digital gene expression matrix was combined with DGEs of *Hormad1*-/- Drop-seq data.

###### Quality Control for Wildtype Single-cell RNA Sequencing Data

DGEs from all wild type experiments were combined into one DGE matrix for further quality control steps. We used a R package, SingleCellExperiment, to create R object that stores gene expression information, dimensionality reduction coordinates, cell-specific size factors for normalization and meta-data. The information for meta-data included experimental batch, type of sample preparation, and type of sample. In addition, the R package scater was used to remove poor quality cells. First, cells were filtered based on the total number of unique molecule identifier(UMI) counts detected per cell. For the wildtype-only analysis, we filtered cells with less than 440 UMI counts. Then, cells with less than 370 unique genes were removed. We also filtered cells based on the ratio between mitochondrial RNAs to total RNAs. This ratio was used to remove cells that are likely being dead or stressed. Cells with greater than 5% of this ratio were removed. Genes that were not detected in at least two cells with more than one transcript count were filtered. After a series of filtering steps, the final DGE had 19153 cells with 11977 genes. This filtered digital expression matrix was used for Seurat downstream analyses. The mean UMI count was 1295 and mean number of expressed genes was 793. We corrected for differences in library sizes of batches using scran, a R package that is developed for library size normalization in single-cell RNA sequencing.

###### K-means clustering and differential expression analysis

After quality control and normalization steps, Seurat was used to perform clustering and visualization of the data. A Seurat R object was created from the post-QC DGE without additional quality control and normalization steps. Highly variable genes were identified and used for performing Principal Component Analysis (PCA) to reduce the dimensionality of the data. A number of statistically significant principle components (PCs) to use for clustering was defined by plotting the variability explained by each PC in decreasing order and determine statistically significant PCs. Cells were clustered by K-nearest neighbor (KNN) graph-based algorithm implemented in Seurat. The resolution of the clustered data will always be either under or over-clustered so to address this issue, we adapted the suggestion by Seurat developers. We first slightly over-clustered data (i. e. created more clusters than necessary) and merged clusters that are transcriptionally indistinguishable. To test for which clusters to be merged, the out-of-bag error (OOBE) method from a random forest classifier was used. We noticed that cells with low number of genes and UMIs were forming clusters on t-SNE plot so these clusters were removed. Finally, differentially expressed markers of all clusters were identified using Seurat and this information was used as an input for generating potential novel cell-type specific markers for validation experiments via immunofluorescence.

###### Sparse Decomposition of Arrays (SDA)

Combined raw DGEs were processed through a series of quality control and normalization steps. Cells were fewer than 200 UMI counts were removed and genes in lower half of expression means were removed. Cells were normalized by square root transformation of total transcript counts per cell and genes were normalized to unit variance. All expression values were capped to maximum of 10. Then, SDA was run with 50 components for 10000 iterations on the filtered and normalized data which had 20322 cells and 19262 genes. Briefly, SDA decomposes a DGE into a number of components represented by two matrices. The columns vectors of the first matrix indicate how much a given component is active in each cell and the rows of the second matrix indicates which genes are active in a given component. SDA convergence was confirmed using the change in free energy, as well as the change in fraction of posterior inclusion probabilities (PIPs) less than 0.5. The distribution of PIPs, cell scores, and gene loadings were also assessed. SDA was also run five times with different seeds as well as with different number of components to ensure stability of the results. Components with a single high loading in one cell (1, 46, 4, 18, 14) were removed to visualize relationships between the components. To visualize and quantify the biological relationships among cells, t-SNE was run on a version of the component scores matrix with potential technical artifacts and batch components removed, using a perplexity of 50. Technical components were manually identified as meeting one or both of the following criteria: two batches of the same mouse line had opposite or very different cell scores (components 6, 12, 22, 28, 41, 29) and or if the highest loading genes were all or mostly ribosomal or pseudogenes (components 25, 9, 43). To assess uncertainty in the t-SNE embedding t-SNE was run multiple times with different seeds (**Figure S6**)

To generate a pseudo-timeline we used a similar approach to that implemented in SCUBA. We iteratively fit a principal curve through the t-SNE plot with increasing degrees of freedom from 4 to 9 using the curve from the previous run as the starting point. Each cell was then assigned to the closest position on this curve. Somatic cells and the Hormad1 X-activated cells were excluded during pseudotime construction. Somatic cells were defined by thresholding the cell scores of somatic components.

Imputed gene expression values (the posterior means of the SDA model) were computed as the matrix product of the cell scores and gene loadings matrix from SDA.

Component names were assigned based on known maker genes from the literature and cross checked for consistency against clustering of the components by t-SNE using the absolute gene loadings or cell scores matrix with a low perplexity (2). Components representing batch effects were identified by plotting cell scores by experimental batch and checking for biological subgroups with opposing cell scores. Plotting the cell scores on t-SNE also revealed ‘patchy' patterning for technical components compared to biological components which tended to vary instead by pseudotime.

We also ensured the KO cells were not unduly affecting the estimated components by separately performing an SDA analysis with only WT cells (normalised separately but with the same parameters). The same number of iterations, number of components, and random seed, were used. We correlated the gene loadings of the Mixed WT & KO SDA analysis with the WT only analysis and found strong correspondence for those WT components which contained many cells **(Figure S7**).

###### Validation of SDA Imputation

In order to formally quantify the accuracy of SDA imputation, we performed a validation study comparing the ability of SDA imputation to correctly predict single cell gene expression data compared to two other simpler approaches. First, we randomly split the post-QC RNA-sequencing reads from the full dataset into two batches: with 20% probability a read is assigned to the “test” dataset, and with 80% probability it is assigned to the “training” dataset. Next we create three predictors of gene expression levels for each cell, using the “training” dataset: we call these “Imputed”, “Cellwise Training” and “Average Training”. The “Imputed” predictor is the SDA-imputed expression level for each gene and each cell based on running SDA on the “training” dataset. The “Cellwise Training” predictor is simply the expression level of each gene in the training dataset for that cell, scaled to the library size of the cell. The “Average Training” predictor for each combination of gene & cell is the average expression level of the gene across all cells in the dataset.

To compare the accuracy of the three predictors for gene expression imputation, we evaluate an objective function for each predictor and each cell, which we call the “quantitative predictive accuracy curve” or QPAC. The QPAC for each predictor is created by rank ordering all genes in a single cell by the predicted level of expression of those genes, from high-to-low (**Figure S5**). For each rank (abscissa), the ordinate is the cumulative sum of “test” data reads for all genes up to that rank (i. e. all genes with higher predicted expression than the current rank). The QPAC is similar in spirit to a receiver operating characteristic (ROC) curve. The area under the curve (AUC) for each QPAC is informative about prediction accuracy; a completely random predictor is expected to produce an AUC of 0.5, while a method with some predictive utility will have an AUC > 0.5. A perfect predictor will have an AUC approaching 1.0 (but the maximum possible AUC will be determined by the true data). A higher level summary of the results was obtained by calculating an average AUC across all cells for each predictor.

###### Transcription Factor Motif Analysis

To discover *de novo* motifs enriched in each component we used the MotifFinder software - an iterative Gibbs sampler described in ^31^. We ran MotifFinder on the repeat masked promoter sequences from Mus Musculus GRCm38 of the top 250 positive and negative genes (separately) for each component. For each component 10 different regions around the TSS were used (150, 200, 250 bp upstream and downstream of the TSS (separately), and 200, 300, 400, and 800 bp centered on the TSS). Each run was repeated three times with different seed motifs (each of length 6bp) and each of those repeated three times with different seeds. MotifFinder was run for 100 iterations in each case.

The resulting *de novo* motifs were annotated with known motifs from the HOCOMOCO database (V11) using Tomtom from the MEME suite. We subset the matches by taking the match with the minimum q-value for each HOCOMOCO target for those with a q-value of less than 0.001. This resulted in 124 different matches from 92 *de novo* motifs. These were manually grouped into 16 categories based on similarity of the motifs. Within each group a single ‘most likely acting' motif was chosen based on external suggestive evidence such as if a knock out of the TF causes infertility, if the TF is specifically expressed in testis by RNA-Seq data, or if the TF is previously known to bind testis expressed genes by ChIP-Seq.

To find good motifs with poor matches to currently known motifs we plotted the sum of the information content by the E value for each *de novo* motif. We found a cluster of motifs with poor E values but large total information content – most of these matched best to ATF1.

In order to determine patterns of association of motifs with pseudotime, we performed correlation tests between the gene loadings of each component and the probability of the motif being present at the promoter of these genes. To find probability of motif presence per gene promoter we used MotifFinder but fixed the position weight matrix to either one of our *de novo* motifs (**Figure 4**) or a motif from the HOCOMOCO database (**Figure S7A**) and ran MotifFinder for 20 iterations. We assessed convergence through the change in proportion of sequences containing the motif.

We used published ChIP-Seq data for the transcription factors *Mybl1 ^32^*, *Crem* ^59^, and *Rfx2* ^33^ to validate our conclusion that some SDA components correspond to genes coregulated by these transcription factors. We performed a Fisher's exact test on the overlap of the genes suggested from the ChIP-Seq studies for each of the three transcription factors with the top 500 genes for each SDA component (positive and negative loadings separately). For *Mybl1* the genes are those that Bolcun-Filas had determined as potential direct targets of MYBL1 (those that were bound in ChIP and were mis-regulated in *repro9* mice, from their Table 1). For *Crem* the genes were those that are “found in the cross-section between DE genes from Kosir et al. and genes bound by CREM in testis from Martianov et al” (i. e. the top 50 genes from Kosir, et al. Supp Table 4). For *Rfx2* the genes are the genes from Kistler et al. that are bound by RFX2 by ChIP-seq and are statistically significantly downregulated at P30 (from their Table S1).

## QUANTIFICATION AND STATISTICAL ANALYSIS

### Apolipoprotein E (ApoE) Immunofluorescence Signal Quantification

To quantify the frequency of ApoE protein signal in wild-type and mutant animals, we counted the total number of intact testicular tubules present on slides and the number of tubules with ApoE protein signal using a confocal microscope at 20x. A Fisher's exact test was used to test the hypothesis that the frequency of ApoE-positive tubules was the same in wildtype, *Mlh3-/-* and *Hormad1-/-* strains.

## DATA AND SOFTWARE AVAILABILITY

Raw data and processed files for Drop-seq and 10X Genomics experiments are available under GEO accession number GEO: GSE113293

R markdown files that enable simulating main steps of the analysis are available upon reasonable request. Custom R code used is available at http://www.github.com/MyersGroup/testisAtlas and archived at DOI: 10.5281/zenodo.1311483.

SDA is available from https://jmarchini.org/sda/

### Supplemental Information

**Table S1**. Summary of all wildtype and mutant single-cell RNA-sequencing experiments.

**Table S2**. Summary of all differentially expressed genes in total joint wildtype and mutant cell clusters

**File S1** A ZIP file containing results of the full SDA analysis reported in the manuscript, which can be loaded and explored in the R computing environment.

